# TWIRLS, an automated topic-wise inference method based on massive literature, suggests a possible mechanism via ACE2 for the pathological changes in the human host after coronavirus infection

**DOI:** 10.1101/2020.02.27.967588

**Authors:** Xiaoyang Ji, Chunming Zhang, Yubo Zhai, Zhonghai Zhang, Chunli Zhang, Yiqing Xue, Guangming Tan, Gang Niu

## Abstract

Faced with the current large-scale public health emergency, collecting, sorting, and analyzing biomedical information related to the “coronavirus” should be done as quickly as possible to gain a global perspective, which is a basic requirement for strengthening epidemic control capacity. However, for human researchers studying the viruses and the hosts, the vast amount of information available cannot be processed effectively and in a timely manner, particularly when the scientific understanding may be limited, which can further lower the information processing efficiency. We present TWIRLS, a method that can automatically acquire, organize, and classify information. Additionally, independent functional data sources can be added to build an inference system using a machine-based approach, which can provide relevant knowledge to help human researchers quickly establish subject cognition and to make more effective decisions. TWIRLS can automatically analyze more than three million words in more than 14,000 literature articles in only 4 hours. Combining with generalized gene interaction databases creates a data interface that can help researchers to further analyze the information. Using the TWIRLS system, we found that an important regulatory factor angiotensin-converting enzyme 2 (ACE2) may be involved in the host pathological changes on binding to the coronavirus after infection. After triggering functional changes in ACE2/AT2R, an imbalance in the steady-state cytokine regulatory axis involving the Renin-Angiotensin System and IP-10 leads to a cytokine storm.

## Introduction

The sudden outbreak of the new coronavirus (SARS-CoV-2) at the end of December 2019 poses a huge threat to human health worldwide. The SARS-CoV-2 virus causes severe respiratory disease (COVID-19) that can quickly spread from person to person and in some cases lead to death. Researchers have found that the new SARS-CoV-2 and SARS coronaviruses invade human cells in target tissues in a similar manner via high-affinity binding to angiotensin-converting enzyme 2 (ACE2)^[1]^. In recent epidemiological investigations of the spread of the SARS-CoV-2 and a preliminary study of the clinical characteristics of this disease^[2-6]^, researchers have found that patients infected with the new coronavirus have severe symptoms similar to those of the SARS infection. The first batch of clinical data reports of SARS-CoV-2 infection cases in China revealed “cytokine storms” in critically ill patients^[7, 8]^. However, the mechanism of the viral infection and pathological changes in the immune system is still lacking. The sooner this information is added to the current clinical knowledge of these viruses, the better the control and treatment of this disease.

Here, we present an automated topic-wise inference method called TWIRLS (<u>T</u>opic-<u>w</u>ise <u>i</u>nfe<u>r</u>ence engine of massive biomedical <u>l</u>iterature<u>s</u>) for processing the massive biomedical literature to summarize coronavirus host-related entities. TWIRLS is capable of collecting, classifying, and analyzing reported coronavirus studies to reveal these entities based on the distribution of specific genes in the text of the articles. By combining with general protein interaction data, links between certain functional cellular/physiological components can be inferred to fill the knowledge gaps on the probable mechanism of host pathological changes. Based on the literature related to the coronavirus, TWIRLS revealed that the altered function of ACE2/AT2R in the host after coronavirus infection possibly leads to an imbalance in the Renin-Angiotensin System (RAS) inducing a cytokine storm. The triggered cytokine storm eventually leads to acute lung injury in the host. Therefore, TWIRLS can be used to guide human researchers by providing further potential therapeutic target information for the treatment of acute viral lung injury based on the regulation of RAS.

## Results

### Coronavirus-study specific entities and host genes

As of February 21, 2020, the PubMed database included 14,878 biomedical articles on coronaviruses. We obtained text data (called local samples) from all related articles on the coronavirus that had been peer reviewed and published by human experts, which included the title, abstracts, author and affiliation information (total 3,182,687 words). The goal of the literature mining was to identify host genes and entities that are relevant to coronavirus research and to establish connections between them. An entity can refers to a word or phrase of the concept name (including related concepts, e.g., virus structure and chemical composition, source of infection, and virus type). The gene names were defined using the mammalian official gene symbols in the Hugo Gene Naming Committee (HGNC) database. We directly retrieved 667 candidate genes from the local samples. By establishing a random distribution of one of the candidate genes in a control sample, the significance of this gene appearing in the local samples can be determined when the frequency of the current gene is an outlier of the random distribution of the control samples (see Methods for details). By calculating the odds ratio, we can also further determine the specificity of the association between this gene and the local samples. In this paper, we selected an odds ratio > 6 as the threshold for this judgment, which resulted in 123 coronavirus study-specific host genes (CSHGs).

To determine the specificity of the entity, we made a choice between different texts in the local samples. We removed numbers, symbols, verbs, and garbled characters to obtain clean versions of the local samples. The coronavirus study-specific entities (CSSE) were then identified in only the clean texts containing CSHGs. Based on the clean selected samples, we next built a local dictionary of candidate CSSEs containing 49,293 words after deduplication. Before calculating the random distribution of each entity, we included the synonymous entities into a same entity number (including singular or plural words, active and passive forms, different tenses, suffixes that do not change the meaning, etc.). For example, synonymous entities such as coronaviral, coronavirus, coronaviruses were grouped into one entity as coronavirus and assigned the same number (see entity number in Table S1, Sheet 1 first column). The previous method of merging synonymous entities was based on a dictionary^[9, 10]^, which not only relied on the integrity of the dictionary, but also required a long retrieval time. To automatically solve the synonymous entity problem, TWIRLS classifies similar strings based on whether there is a significant statistical association between the character blocks in a set of candidate entities including various synonymous entities (see Methods). After cleaning and processing, CSSEs were identified by TWIRLS using a similar method to that for CSHG as described above.

For the candidate CSSE dictionary, a random distribution model for each entity was built by TWIRLS using the control samples. We identified 623 CSSEs (Table Sl, Sheet 1) based on the outliers discriminated by the random model and calculated odds ratio. For example, TWIRLS found 100 CSSEs close to ACE2, the receptor of SARS and SARS-CoV-2 viruses (see left panel in Figure 1). The size of the entity represents the relative distance to ACE2, with a larger size indicating a closer distance to ACE2. Additionally, we present the CSSE cloud of the human receptor gene DPP4 of the MERS virus (see right panel in Figure 1).

**Figure 1.**
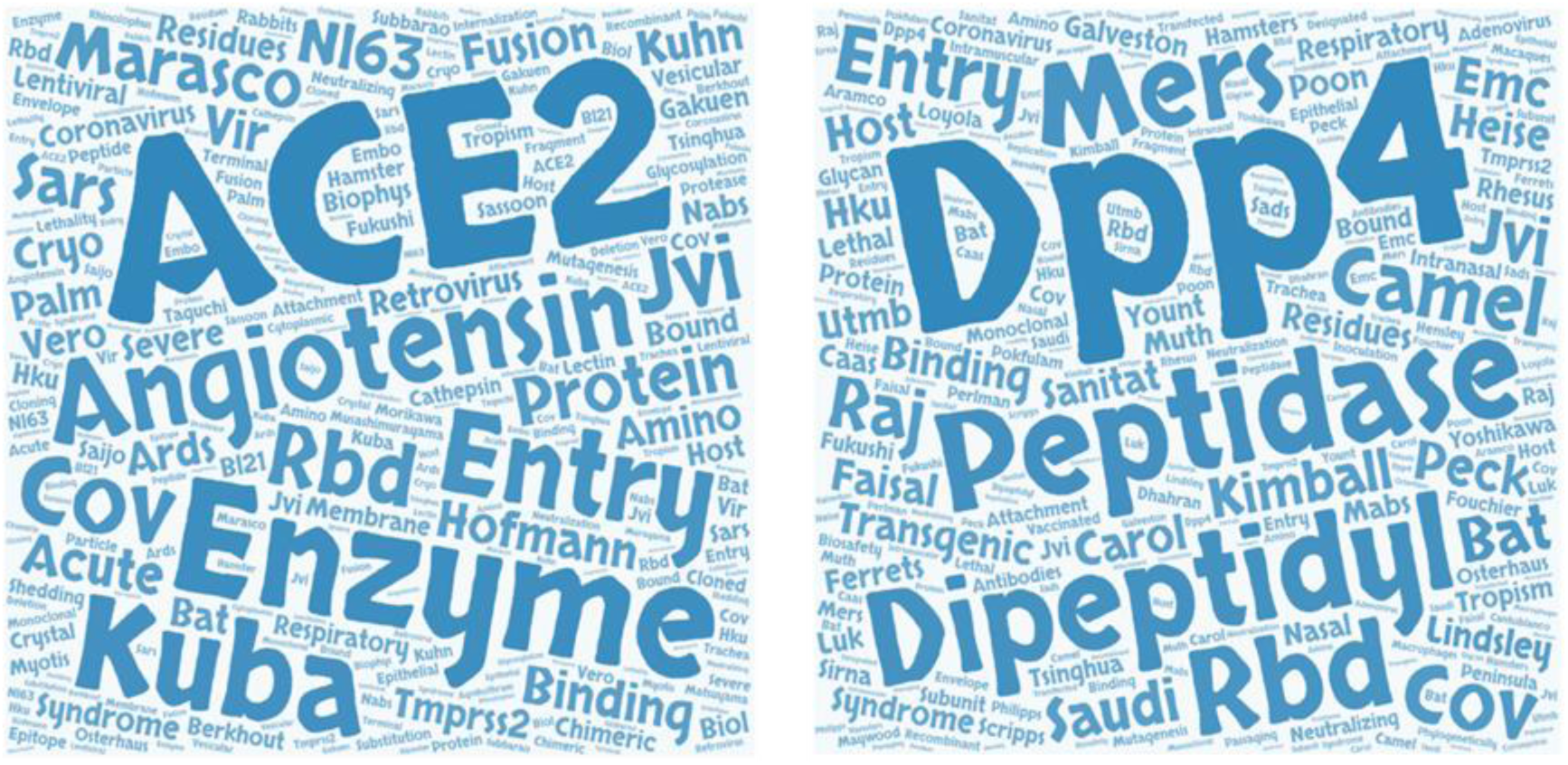
The entity cloud (CSSE cloud) associated with ACE2 and DPP4 in the coronavirus knowledge graph.

### Categories of entity and their labels-human conclusion and enriched pathway

Although TWIRLS only identified 623 CSSEs after collation, for human researchers, the information is scattered in words, which is limited for reconstructing understandable mechanistic models. Therefore, TWIRLS clusters CSSEs according to the rules defined by CSHG distribution, as genetic level research can accurately answer and solve physiological and pathological problems. TWIRLS first calculated the specific co-distribution between CSHGs in local samples, then determined the distance between each pair of CSSEs and performs dichotomy clustering according to the linkage relationship between CSSEs and CSHGs. This classified the 623 entities into 32 categories represented as C0-C31 (see category number in Table S1, Sheet 1 second column). In addition, for each category, TWIRLS also cited the top ten most relevant references for human researchers (Table S2). Therefore, in any category, according to the CSSE and the most relevant literature, we can quickly provide “Labels of conclusion-drawn-by-human-researcher” (HR Labels) for this category. This label outlines the most relevant research directions of the current entity category. For example, for category C3, the HR label is “Neurotrophic Coronavirus Related to Immune-Mediated Demyelination”. We have summarized the HR labels for the 32 entity categories in Table 1.

**Table 1.** Coronavirus-entity category labels and genes associated with each category. MISC indicates the label cannot be summarized.

| Category | HR label |
| --- | --- |
| C0 | MISC |
| C1 | Canine coronavirus |
| C2 | Porcine epidemic diarrhea (PED) |
| C3 | Neurotropic coronavirus correlated with immune-mediated demyelination |
| C4 | Coronavirus that infects humans |
| C5 | Coronavirus spike protein |
| C6 | Protease enhances SARS-CoV infection |
| C7 | Monoclonal antibody to the coronavirus nucleocapsid protein |
| C8 | SARS-CoV genome |
| C9 | Avian infectious bronchitis coronavirus |
| C10 | Coronavirus and interferon |
| C11 | Feline infectious peritonitis (FIP) |
| C12 | Vectors of novel coronaviruses |
| C13 | Mouse hepatitis virus |
| C14 | Interaction between coronaviruses and receptors |
| C15 | Coronavirus-related vaccines |
| C16 | Identification of MHC class I restricted T-cell epitopes |
| C17 | Transmissible gastroenteritis coronavirus |
| C18 | SARS coronavirus inhibitors and diagnostic methods |
| C19 | Coronavirus fusion with host cells and virus replication |
| C20 | Gene Therapy-Inhibition of Coronavirus by Antisense RNA, Sense RNA and Protein |
| C21 | Clinic Imaging |
| C22 | Cytotoxic T-lymphocyte escape |
| C23 | SARS coronavirus compound inhibitors |
| C24 | Coronavirus studies using biophysical methods |
| C25 | Detection of viral pathogenicity and distribution (RT-PCR, immunohistochemistry and in situ hybridization) |
| C26 | MISC |
| C27 | MISC |
| C28 | Effects of coronavirus infection on the body |
| C29 | MISC |
| C30 | Human respiratory coronavirus NL63 |
| C31 | MISC |

The relative position of any CSHG to a certain CSSE can be estimated by TWIRLS (see Table S1, the ranking matrix in Sheet 1). As each category contains different entities, we can determine whether a certain CSHG is significantly closer to each entity in the current category based on the ranking matrix between CSHG and CSSE. For example, the average distance between ACE2 and the 92 entities in category C5 is first calculated, then a random distribution model of the average distance between ACE2 and any of the 92 entities (3000-5000 times) is built, and finally, we determine if the average distance between ACE2 and entities in category C5 is significantly less than and deviates from the mean of the random distribution (Z score = -5.8416). The significance of each category associated with each CSHG is then scored by TWIRLS ranging between -10 and +10, with a smaller score indicating the current CSHG is more relevant to the current category (see the Z-score matrix in Table S1, sheet 2). For an entity category, the associated CSHGs (e.g., ^Ci^CSHGs, where *i* represents the category number) can thus be selected by a Z score <-3 (the Z scores describing the association between CSHG and any category is summarized in Sheet2 of Table S1, and the category labels of all CSHGs are provided in Sheet 3).

Specifically, Spike proteins (S proteins) of different coronaviruses recognize different receptor molecules on human cells, such as ACE2 (binds to Spike proteins in SARS and SARS-CoV-2 virus) and DPP4 (binds to Spike protein in MERS virus). We found that these two genes are assigned to the C5 category, which has a corresponding HR label of “Spike protein (S) of coronavirus”, suggesting that TWIRLS can automatically provide an interface to summarize human findings and help human experts quickly understand the research directions and necessary knowledge in this field.

The distribution and meaning of the data can be compared to specific expression values of CSHG under different conditions (here, the category is used as a condition). Therefore, based on the distribution of the pathway signatures, TWIRLS can recommend the most likely and least likely signaling pathways (Table 2). On the other hand, TWIRLS can also recommend the most likely and least likely categories for each signaling pathway. As an example, Table 3 shows the signaling pathways most likely associated with category C3 with the most unlikely corresponding category.

**Table 2.** The most relevant and least relevant signaling pathways of each coronavirus-entity category

| Class | Likely pathway | Z score | Unlikely pathway | Z score |
| --- | --- | --- | --- | --- |
| C0 | PKCθ Signaling in T Lymphocytes | 1.5782 | Toll-like Receptor Signaling | -1.7195 |
| C1 | AMPK Signaling | 5.1816 | TGF-β Signaling | -4.0841 |
| C2 | Extrinsic Prothrombin Activation Pathway | 4.3314 | Melanocyte Development and Pigmentation Signaling | -3.8316 |
| C3 | Renin-Angiotensin Signaling | 5.6100 | AMPK Signaling | -4.4763 |
| C4 | April Mediated Signaling | 3.2382 | Neuregulin Signaling | -2.4121 |
| C5 | Melanocyte Development and Pigmentation Signaling | 3.7887 | Extrinsic Prothrombin Activation Pathway | -3.9823 |
| C6 | Leukocyte Extravasation Signaling | 2.3615 | Pancreatic Adenocarcinoma Signaling | -2.3694 |
| C7 | Role of BRCA1 in DNA Damage Response | 4.6871 | NF-κB Signaling | -4.8545 |
| C8 | PKCθ Signaling in T Lymphocytes | 1.9902 | VDR_RXR Activation | -2.6143 |
| C9 | Toll-like Receptor Signaling | 4.2000 | Role of BRCA1 in DNA Damage Response | -2.5017 |
| C10 | April Mediated Signaling | 4.3113 | TGF-β Signaling | -3.5853 |
| C11 | Acute Phase Response Signaling | 2.5066 | Renin-Angiotensin Signaling | -3.2492 |
| C12 | ATM Signaling | 2.7020 | PKCθ Signaling in T Lymphocytes | -1.9846 |
| C13 | Retinoic acid Mediated Apoptosis Signaling | 2.6069 | Interferon Signaling | -1.8953 |
| C14 | ATM Signaling | 2.9244 | PKCθ Signaling in T Lymphocytes | -1.9667 |
| C15 | AMPK Signaling | 1.9142 | Leukocyte Extravasation Signaling | -2.4233 |
| C16 | Role of BRCA1 in DNA Damage Response | 3.1033 | Colorectal Cancer Metastasis Signaling | -3.1033 |
| C17 | Production of Nitric Oxide and Reactive Oxygen Species in Macrophages | 3.1398 | mTOR Signaling | -2.8893 |
| C18 | Extrinsic Prothrombin Activation Pathway | 1.8215 | Intrinsic Prothrombin Activation Pathway | -1.5680 |
| C19 | Role of NFAT in Regulation of the Immune Response | 2.0089 | CD40 Signaling | -2.2696 |
| C20 | April Mediated Signaling | 2.3687 | Aryl Hydrocarbon Receptor Signaling | -1.5471 |
| C21 | NRF2-mediated Oxidative Stress Response | 1.4679 | CD40 Signaling | -2.4042 |
| C22 | Toll-like Receptor Signaling | 2.2825 | Melanocyte Development and Pigmentation Signaling | -1.6007 |
| C23 | ATM Signaling | 2.5518 | mTOR Signaling | -1.8228 |
| C24 | Melanocyte Development and Pigmentation Signaling | 2.1700 | Aryl Hydrocarbon Receptor Signaling | -2.1700 |
| C25 | Extrinsic Prothrombin Activation Pathway | 1.9825 | IL-22 Signaling | -1.7638 |
| C26 | mTOR Signaling | 2.7449 | Aryl Hydrocarbon Receptor Signaling | -1.7790 |
| C27 | AMPK Signaling | 2.4033 | Intrinsic Prothrombin Activation Pathway | -1.7579 |
| C28 | NRF2-mediated Oxidative Stress Response | 2.0846 | PKCθ Signaling in T Lymphocytes | -1.3415 |
| C29 | TGF-β Signaling | 2.4379 | Ephrin A Signaling | -2.0030 |
| C30 | Role of NFAT in Regulation of the Immune Response | 2.1383 | Leukocyte Extravasation Signaling | -1.8524 |
| C31 | Role of NFAT in Regulation of the Immune Response | 1.7979 | Neuregulin Signaling | -1.4467 |

**Table 3.**
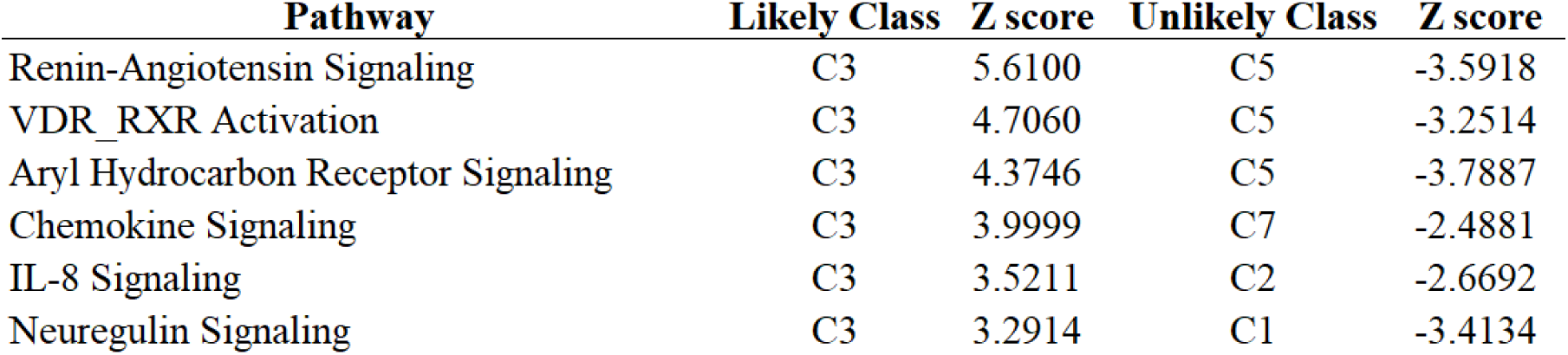
Recommended signaling pathway most relevant to entity category C3

| **Pathway** | **Likely Class** | **Z score** | **Unlikely Class** | **Z score** |
| --- | --- | --- | --- | --- |
| Renin-Angiotensin Signaling | C3 | 5.6100 | C5 | -3.5918 |
| VDR_RXR Activation | C3 | 4.7060 | C5 | -3.2514 |
| Aryl Hydrocarbon Receptor Signaling | C3 | 4.3746 | C5 | -3.7887 |
| Chemokine Signaling | C3 | 3.9999 | C7 | -2.4881 |
| IL-8 Signaling | C3 | 3.5211 | C2 | -2.6692 |
| Neuregulin Signaling | C3 | 3.2914 | C1 | -3.4134 |

### Entity category associated genes involved in generalized interaction networks

We coupled the data with gene interaction/regulation databases and constructed a generalized protein-protein interaction network (PPI network) among 119 genes out of the 123 CSHGs. We defined the direct interaction between two genes as a 1 degree (1°) interaction, and the indirect interaction connecting two genes through a gene as a 2 degree (2°) interaction. All the genes in the 1° networks mined in the PPI database are shown in Figure 2. The results after deduplication showed 2,004 pairs in the 119 CSHGs (see Table S1, Sheet 4). As a control, the average interactions of 119 randomly selected genes in the database showed between 252 to 612 pairs (average 220.16, standard deviation 35.15). Compared to random genes, the regulatory connections between CSHGs were significantly enriched (Z score = 50.97).

**Figure 2.**
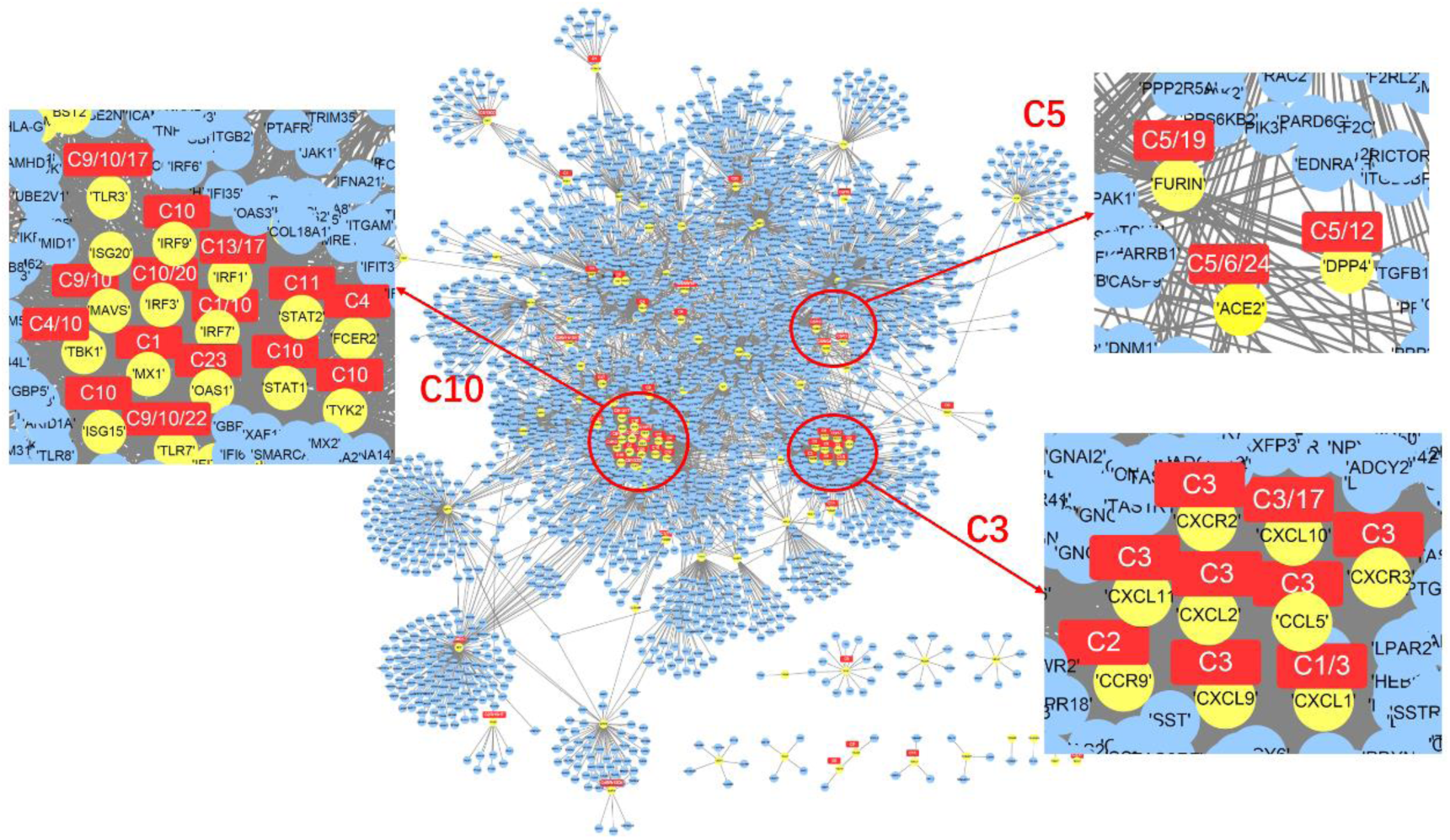
Gene interaction network centered on 119 CSHGs. The yellow nodes represent 119 CSHGs, the blue nodes represent genes that interact with CSHG in the string database (combination score> 800), and the red squares mark the most relevant entity category of CSHG.

Those CSHGs associated with a certain category had much closer interactions. For example, CSHGs associated with category C3 (or associated with C5 or C10) were closer to each other in the 1° networks (Figure 2), suggesting that TWIRLS can possibly highlight important research directions and biology systems involved in coronavirus-specific research and can provide reliable interfaces for further automatic inference.

Several hub genes among the 119 CSHGs were further recommended by TWIRLS. Compared to a random sampling from all interactions recorded in the database, these hub genes had significantly increased numbers of interactions with the other 118 CSHGs. The recommended results showed that the three members of the IFITMs family (IFITM1-3) ranked first, second, and sixth among the top ten hub genes (CSSE cloud of the IFITMs family genes is shown in Figure 3; detailed ranking recommendation results are shown in Table S1, Sheet 5). These IFITMs genes showed 115 interactions, accounting for 8.59% out of all 1,338 interactions of the 119 CSHGs. These IFITMs were significantly enriched in the local samples representing updated coronavirus-related studies (average 0.03% in the control test of random samplings, p <1.5676e-61). The IFITMs family plays crucial roles in the induction of interferons during viral infections. Under the action of interferon, IFITMs disrupts intracellular cholesterol homeostasis and prevents the virus from entering the host cell^[11]^. However, TWIRLS did not directly associate IFITMs with any category, so we needed to provide more information so that TWIRLS can determine which part of these genes might be involved in the coronavirus infection and host body response.

**Figure 3.**
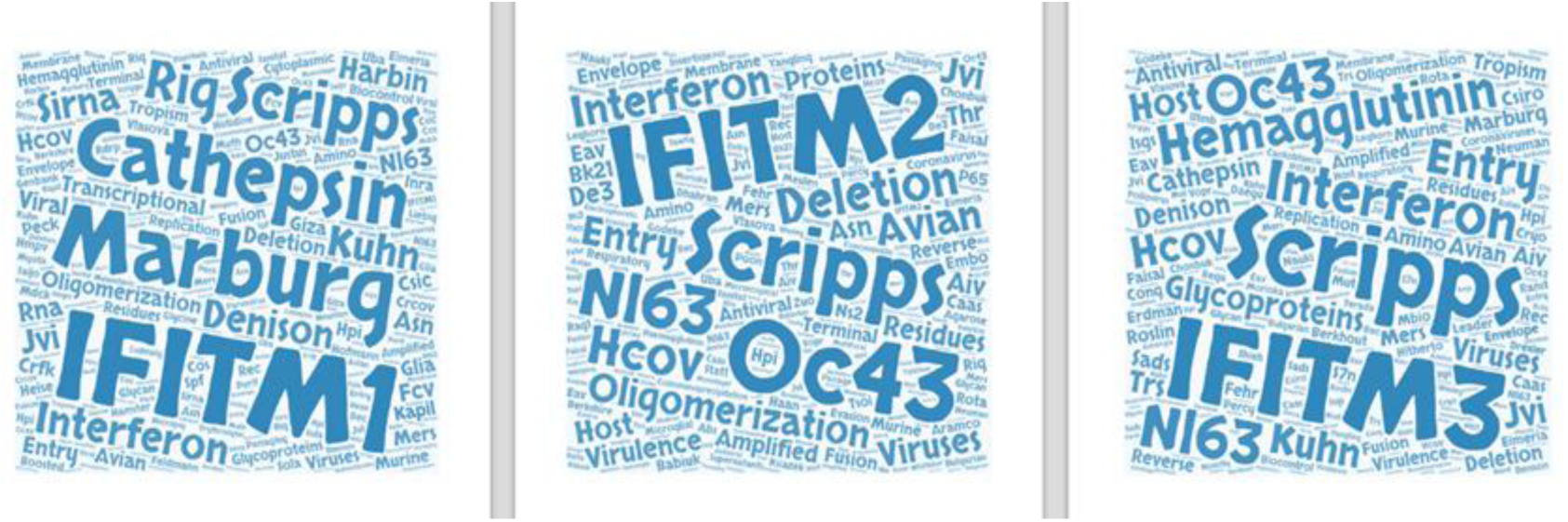
The entity clouds of the three IFITMs family proteins (IFITM1-3) in the coronavirus knowledge graph.

Combining with generalized interaction databases provides richer interactions and regulatory linkages. We extended the 119 CSHGs to their 2° networks based on the interactions with higher likelihood of connections (Combined score> 800). The 2° networks expanded the number of genes from 119 host genes to 3,494 genes that may be associated with coronavirus (see Table S1, Sheet 6 for a list of genes, excluding CS119, as this type of gene is called CSHG2). These genes are mainly involved in two types of functions, virus-related signaling pathways and immune function-related pathways. Table 4 shows a summary of KEGG signaling pathways.

**Table 4.**
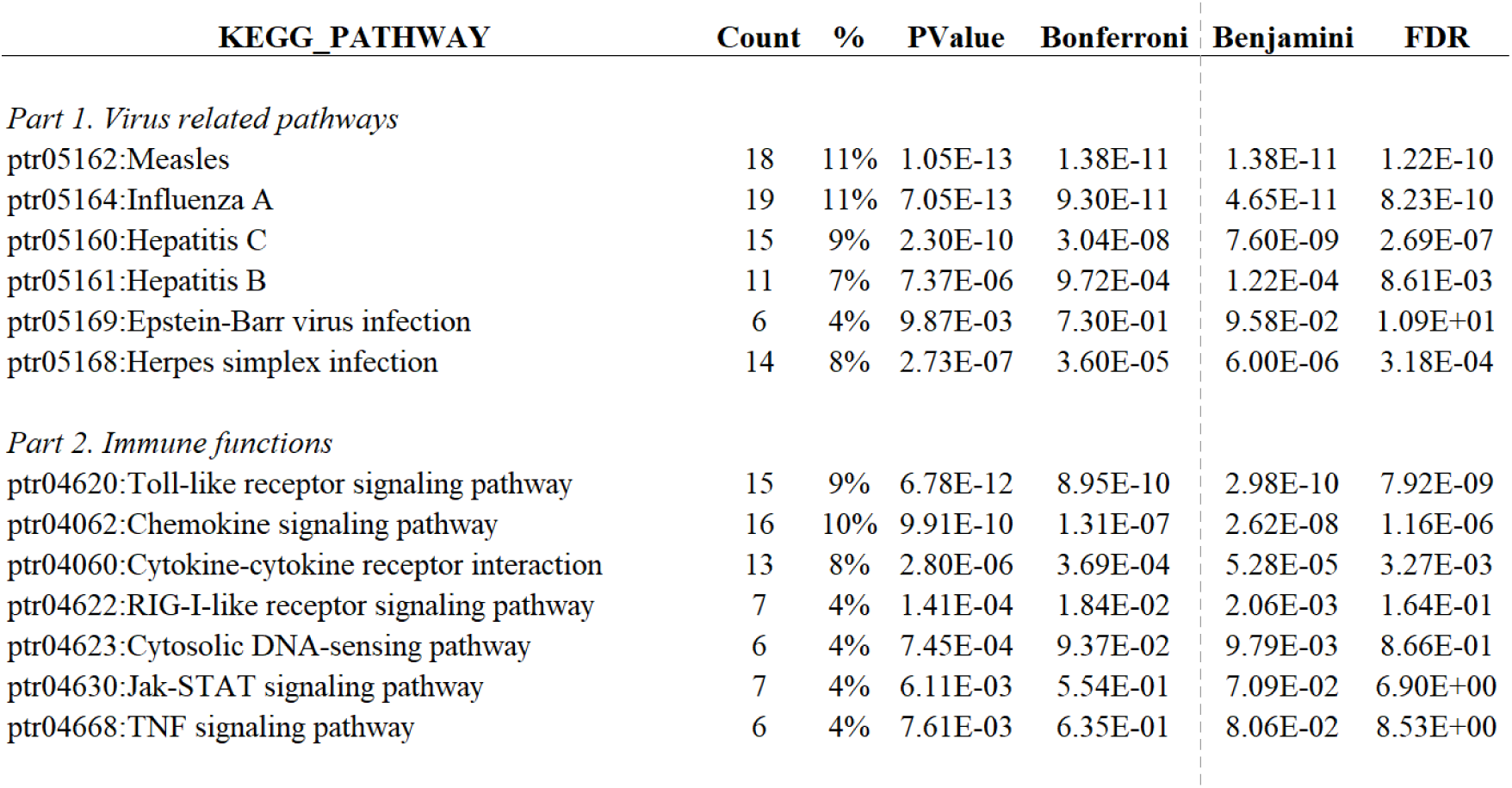
The signaling pathways enriched by 119 CSHGs.

| KEGG_PATHWAY | Count | % | PValue | Bonferroni | Benjamini | FDR |
| --- | --- | --- | --- | --- | --- | --- |
| *Part 1. Virus related pathways* |  |  |  |  |  |  |
| ptr05162:Measles | 18 | 11% | 1.05E-13 | 1.38E-11 | 1.38E-11 | 1.22E-10 |
| ptr05164:Influenza A | 19 | 11% | 7.05E-13 | 9.30E-11 | 4.65E-11 | 8.23E-10 |
| ptr05160:Hepatitis C | 15 | 9% | 2.30E-10 | 3.04E-08 | 7.60E-09 | 2.69E-07 |
| ptr05161:Hepatitis B | 11 | 7% | 7.37E-06 | 9.72E-04 | 1.22E-04 | 8.61E-03 |
| ptr05169:Epstein-Barr virus infection | 6 | 4% | 9.87E-03 | 7.30E-01 | 9.58E-02 | 1.09E+01 |
| ptr05168:Herpes simplex infection | 14 | 8% | 2.73E-07 | 3.60E-05 | 6.00E-06 | 3.18E-04 |
| *Part 2. Immune functions* |  |  |  |  |  |  |
| ptr04620:Toll-like receptor signaling pathway | 15 | 9% | 6.78E-12 | 8.95E-10 | 2.98E-10 | 7.92E-09 |
| ptr04062:Chemokine signaling pathway | 16 | 10% | 9.91E-10 | 1.31E-07 | 2.62E-08 | 1.16E-06 |
| ptr04060:Cytokine-cytokine receptor interaction | 13 | 8% | 2.80E-06 | 3.69E-04 | 5.28E-05 | 3.27E-03 |
| ptr04622:RIG-I-like receptor signaling pathway | 7 | 4% | 1.41E-04 | 1.84E-02 | 2.06E-03 | 1.64E-01 |
| ptr04623:Cytosolic DNA-sensing pathway | 6 | 4% | 7.45E-04 | 9.37E-02 | 9.79E-03 | 8.66E-01 |
| ptr04630:Jak-STAT signaling pathway | 7 | 4% | 6.11E-03 | 5.54E-01 | 7.09E-02 | 6.90E+00 |
| ptr04668:TNF signaling pathway | 6 | 4% | 7.61E-03 | 6.35E-01 | 8.06E-02 | 8.53E+00 |

Among the entire network, we found several CSHGs in the 1° networks (32.6%-35.71%) that directly interacted with three members of the IFITMs family, whereas fewer CSHGs in the 2° network (5.21%-9.46%) indirectly interacted with them. Although there was a higher proportion of directly interacting CSHGs, they were not significantly enriched in any category (see Table S1, Sheet 7 for enrichment scores of 1° network nodes in different categories), whereas the indirect CSHGs were significantly enriched mainly in the C3 and C10 categories (Z score > 3) (see Table S1, Sheet 8 for enrichment scores of 2° network nodes in different categories). These findings demonstrate that TWIRLS can provide new insights about hub molecules, particularly when coupled with interaction information. The new candidate genes, IFITMs, had potential functions associated with category C3, but when adding generalized interaction information, TWIRLS also inferred possible functions of the proteins not associated with any category.

### Reconstruction of mechanic consequence of coronavirus invasion

It is generally considered that the SARS virus binds to ACE2 receptors leading to respiratory tract infections in humans, whereas the MERS virus infects the lower respiratory tract through binding with DPP4^[12]^. The different distribution of these receptors in the respiratory tract results in different degrees of infection. Although the infection ability of MERS is lower than in SARS, the mortality is higher (in about one-third of patients) because of the deeper infection site^[13]^. Similar to the SARS virus, viral genomics and structural biology studies have shown that ACE2 is also a functional receptor for the new SARS-CoV-2 coronavirus. After binding to ACE2 via its Spike protein, SARS-CoV-2 undergoes membrane fusion and enters the host cells by endocytosis. The ACE2 peptidase is a key regulator of the Renin-Angiotensin System (RAS). It is highly expressed in the heart, kidney, and testis, and is also expressed at lower levels in other tissues (mainly in the intestine and lungs)^[14, 15]^. Recent studies have shown that the binding of the S protein to ACE2 in the new coronavirus is 10 to 20 times stronger than in the SARS virus ^[16]^, which may help the new coronavirus infect the host through the upper respiratory tract, significantly increasing its infectivity. Using TWIRLS, we were able to identify both ACE2 and DPP4 genes as CSHGs, and both were significantly associated with the C5 category. The HR label for this category is “associated with S protein.”

In addition to ACE2 and DPP4, other CSHGs that are significantly associated with the C5 category include FURIN and TMPRSS2. The former may be required for the H7N1 and H5N1 influenza virus infections, probably via hemagglutinin-induced lysis, whereas the latter is widely reported to mediate and assist in the invasion of host cells by multiple viruses. Transmembrane protease serine 2 (TMPRSS2) is a serine protease that hydrolyzes and activates the spike glycoproteins of human coronavirus 229E (HCoV-229E), human coronavirus EMC (HCoV-EMC), Sendai virus (SeV) and human interstitial pneumovirus (HMPV), and 1,2,3 fusion glycoproteins of F0, 4a, and 4b human parainfluenza viruses (HPIV)^[17, 18]^. The function of this gene is essential for the transmission and pathogenesis of influenza A viruses (H1N1, H3N2 and H7N9 strains). It is also involved in the hydrolysis and activation of hemagglutinin proteins, which are essential for viral infectivity^[19, 20]^. Although entities in the C5 category and in the cited literature mainly show that virus invasion is facilitated by virus-binding receptors and membrane proteases, the biological mechanism of the receptor binding to viruses leading to pathological changes has been reported less frequently.

TWIRLS recommend new genes that interact with ^C5^CSHGs, and other 1° or 2° CSHGs linked to this gene might be enriched in other categories. These inferences are based on a process that finds new genes connected to different categories. The connected categories suggested potential regulatory relationships between different biological functions or phenotypes. The genes that serve as linkers are potential targets for gain- and loss-of-function experiments to identify those systems described by the meaningful entities in these categories.

In this paper, TWIRLS found the 2° networks showed connections with certain CSHGs associated with categories or with no category. For example, TWIRLS found that CSHGs in the 2° connections of IFITM1 were mainly concentrated in the C3 category (see Figure 4). Interestingly, CSHGs in the 2° connections of ACE2 and DPP4 associated with C5 category were also enriched in C3 category, inferring that the information summarized in C3 category probably describe the underlying mechanisms of the pathological changes after coronavirus infection. In our analysis, the signaling pathways in C3 were mainly RAS, Vitamin D and RXR activation, and Chemokine signaling, with RAS being the most significant (as shown in Table 3, which summarizes C3-related signaling pathways).

**Figure 4.**
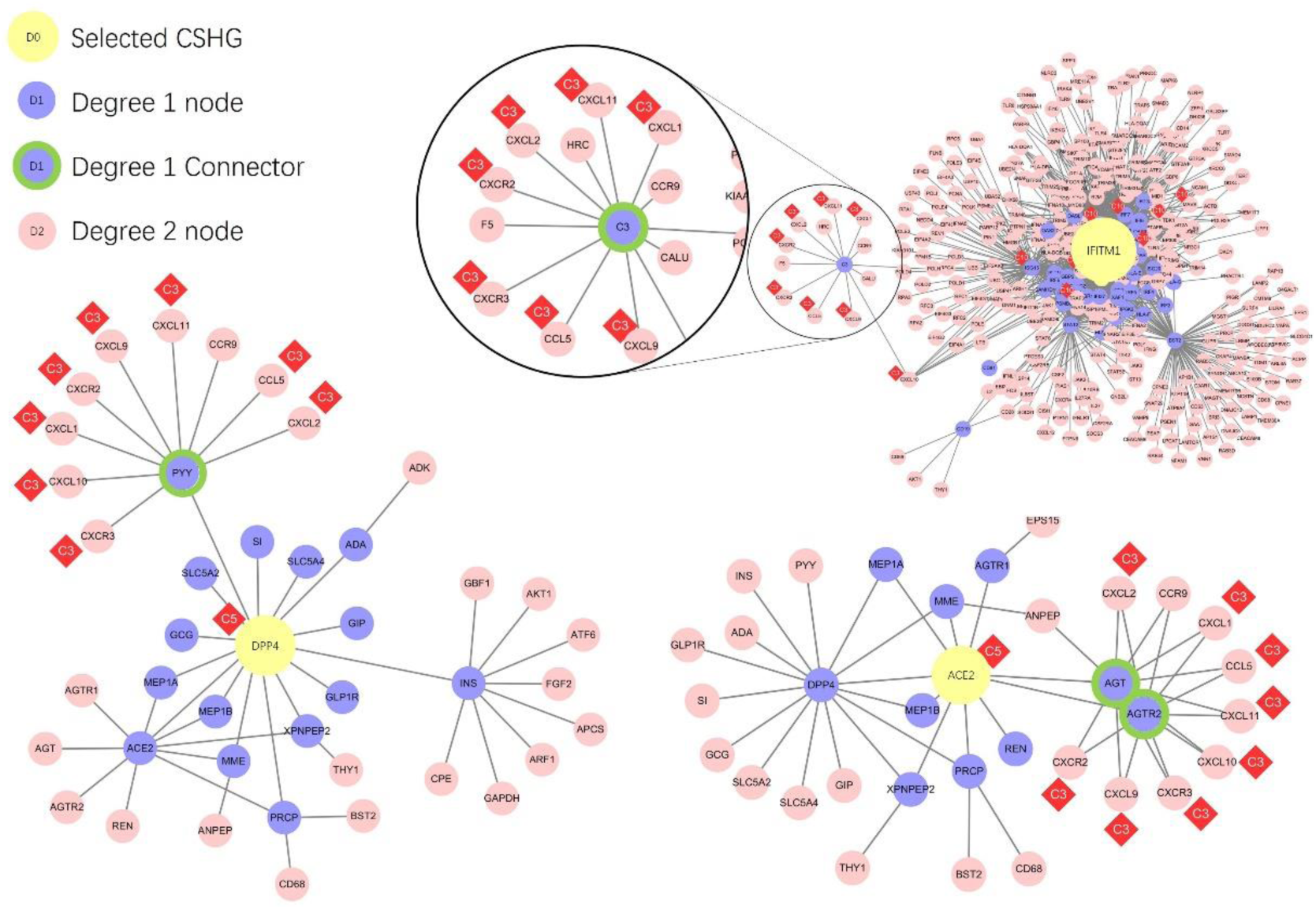
The gene interaction networks centered around DPP4, ACE2, and IFITM1, respectively. The yellow nodes represent the ACE2, DPP4 and IFITM1 genes, purple nodes represent genes that have 1 degree of interaction with the core genes, green circled purple nodes represent the genes connecting CSHG and C3 category-related genes, and pink nodes represent genes with 2 degrees of interaction with the core gene. The red diamonds show the most relevant entity category symbol for CSHG.

Figure 4 shows the CSHGs in the 2° connections of IFITM1, ACE2, and DPP4 were enriched in C3 category through different genes (AGT/AGTR2 in ACE2, PYY in DPP4, and C3 in IFITM1), which then linked to C3-associated cytokines including CCL5, CXCL1, CXCL10, CXCL11, CXCL2, CXCL9, CXCR2, and CXCR3 (Figure 5). Subsequently, these linker genes may contain information on the biological mechanisms that may be important for understanding the disease. For example, TWIRLS recommended angiotensinogen (AGT) and angiotensin II receptor type 2 (AGTR2 or AT2R) genes in the C3 category associated with ACE2. This supports that RAS is probably involved in the pathological changes caused by cytokine storms after S protein binds to ACE2, as suggested by other reports.

**Figure 5.**
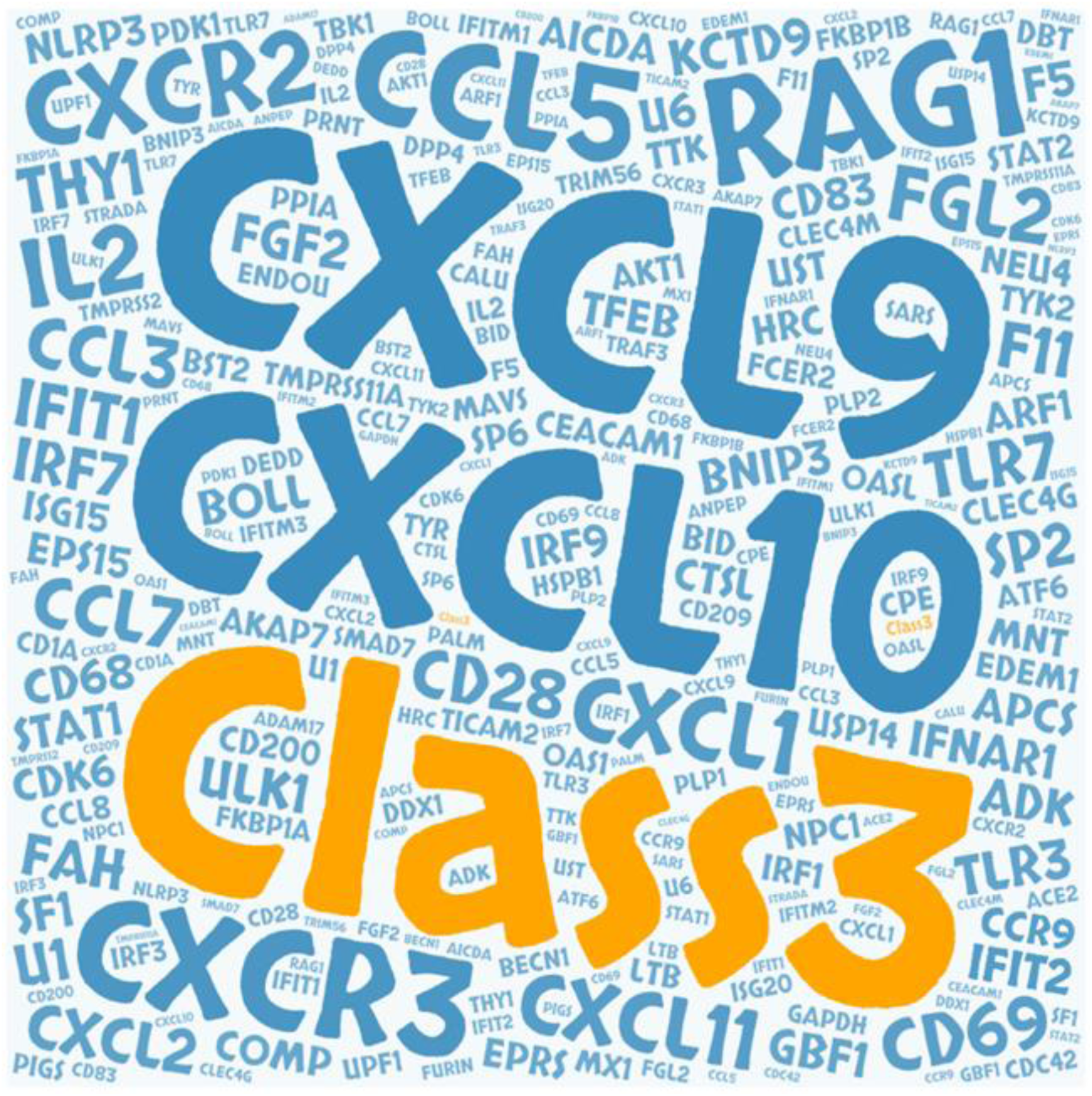
The gene cloud associated with coronavirus-C3 entity category

We next used TWIRLS to calculate the 1° and 2° networks of all 119 CSHGs. Based on the significantly enriched categories of CSHGs in the above networks, TWIRLS separately constructed models for the complex relationships of each CSHG. We found that 45.53% of the CSHGs in these networks were associated with C3 or C10 categories, and five genes (CCL3, CCL5, CXCL1, CXCL2, and STAT2) were associated with both. This suggests that the biological mechanisms described by the C3 and C10 categories might be universally involved. Research on the entities, genes, pathways, and linker genes participating in the C3 and C10 categories could lead to new directions for the prevention, treatment, and clinical management of coronavirus infections.

## Discussion

In this study, we used TWIRLS, a machine-based approach to collect, summarize, and analyze about 15,000 biomedical articles related to coronavirus, with the aim to elucidate the mechanisms underlying coronavirus-induced host pathological changes. Using TWIRLS, we found a possible mechanism involving ACE2/AT2R-RAS-Cytokine signaling, which becomes imbalanced under virus infection leading to cytokine storms. The TWIRLS system is an automated process that can summarize the entities and genes specifically related to coronaviruses. By combining this system with generalized interaction databases, we can reveal further associations that can provide a deeper understanding of the biological mechanisms of the disease phenotype caused by virus-host interactions.

The Renin-Angiotensin system consists of an enzymatic cascade beginning with liver-mediated production of AGT^[21]^. Angiotensin-converting enzyme (ACE), part of RAS, regulates many physiological processes including inflammation and brain functions^[22]^. Angiotensin II (Ang II) is the main effector of this system and is formed by successive enzymatic actions of renin and ACE. It exerts most of its actions through the activation of Ang II type 1 and type 2 receptors (AT1R and AT2R)^[23]^. Deficiency of ACE2 causes respiratory failure pathologies such as sepsis, pneumonia, and SARS^[24, 25]^. It has been confirmed that genetic deletion of AT1a receptor expression can significantly improve lung function and reduce the formation of pulmonary edema compared with wild-type mice^[26]^. In contrast, inactivation of AT2R in mice aggravated acute lung injury. This suggests that AT1R mediates the pathogenicity of Ang II, whereas activated AT2R has a protective role^[27]^. Thus, ACE/AT1R and ACE2/AT2R negatively feedback to one another, playing important roles in RAS-mediated central nervous system and cardiovascular functions. The binding of the virus to ACE2 may disrupt this balance, which causes a steady-state imbalance of RAS, leading to subsequent pathological changes.

Although Ang II was originally described as an effective vasoconstrictor, there is growing evidence that it is closely involved in the inflammatory response of the immune system. Pro-inflammatory cytokines derived from immune cells normally regulate the RAS component, which further accelerates the formation of systemic and local Ang II^[28-30]^. In particular, pro-inflammatory cytokines regulate the production of AGT in the liver and kidney^[31-33]^. On the other hand, RAS has also been implicated in mediating the cytokine storm and has functional relationships with the immune system. Angiotensin II regulates vascular tension and stimulates the release of pro-inflammatory cytokines^[34, 35]^. The production and release of CXC chemokines can induce the accumulation of neutrophils in vivo^[36]^. Meanwhile, ACE inhibitors and Ang II receptor blockers have been used in a number of cytokine-mediated inflammatory pathologies, and AT1R blockers (angiotensin receptor blocker) were shown to have beneficial effects that were commonly attributed to AT2R activation^[37]^. At the same time, it was reported that Ang II-stimulated human endothelial cells had increased release of a CXC chemokine, IP-10. The IFN-γ-inducible protein 10 (IP-10 or CXCL10) is mainly expressed in the lung and is a chemoattractant for activated T cells. The expression of IP-10 has been observed in many Th1-type inflammatory diseases, where it is thought to play an important role in recruiting activated T cells to sites of tissue inflammation. Therefore, RAS dysfunction may result in the accumulation of cytokines, such as in the lungs leading to excessive accumulation of immune cells and interstitial fluid, blocking the airways and causing eventual death. As reported in the first severely infected patients diagnosed with COVID-19, a large number of patients experienced “cytokine storms” that was fatal^[7]^. Figure 6 summarizes the functional changes and pathological consequences of RAS system after ACE2 combines with the coronavirus.

**Figure 6.**
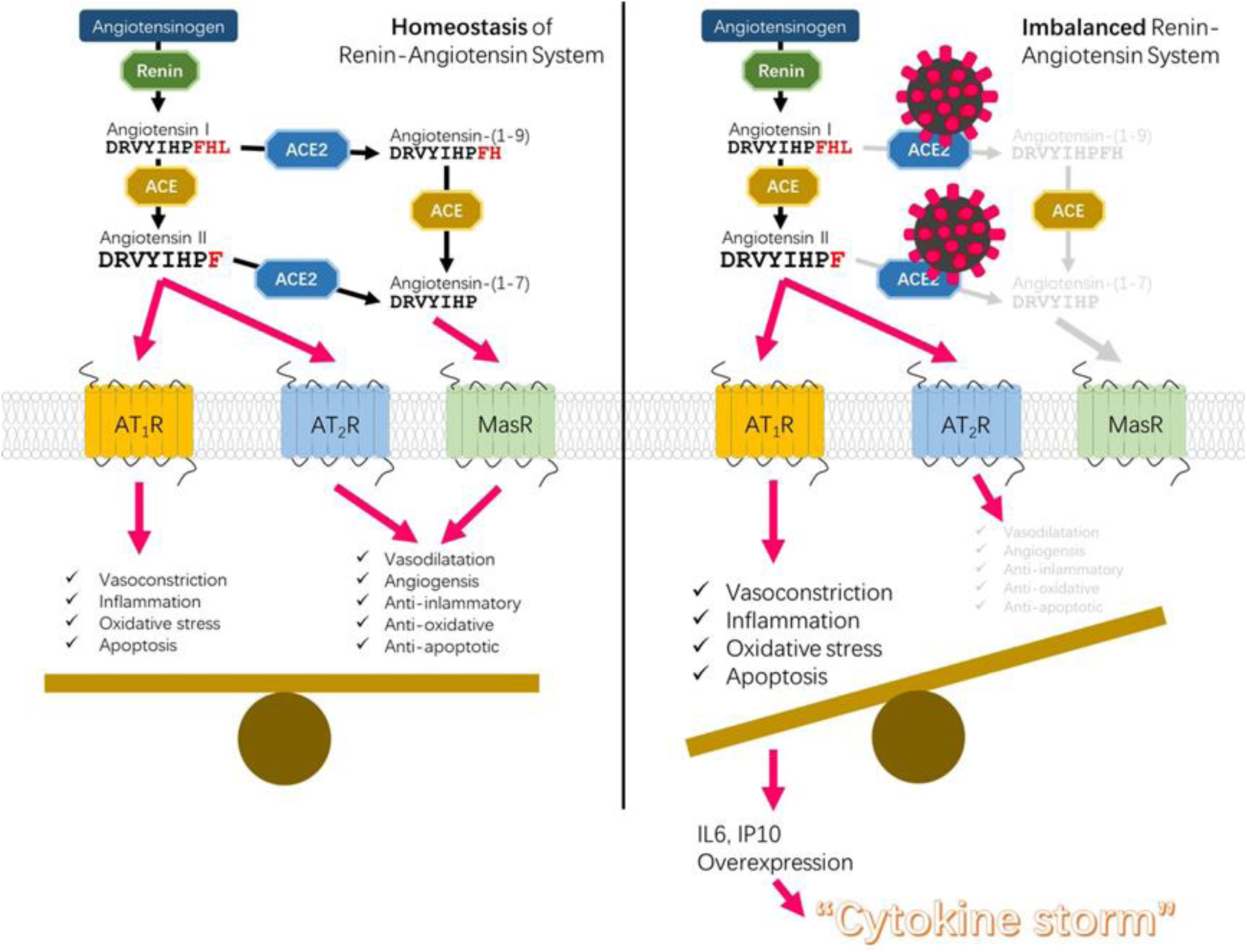
Disequilibrium of RAS-cytokine signaling homeostasis causing cytokine storms triggered by ACE2-mediated coronaviral infection

**Figure 7.**
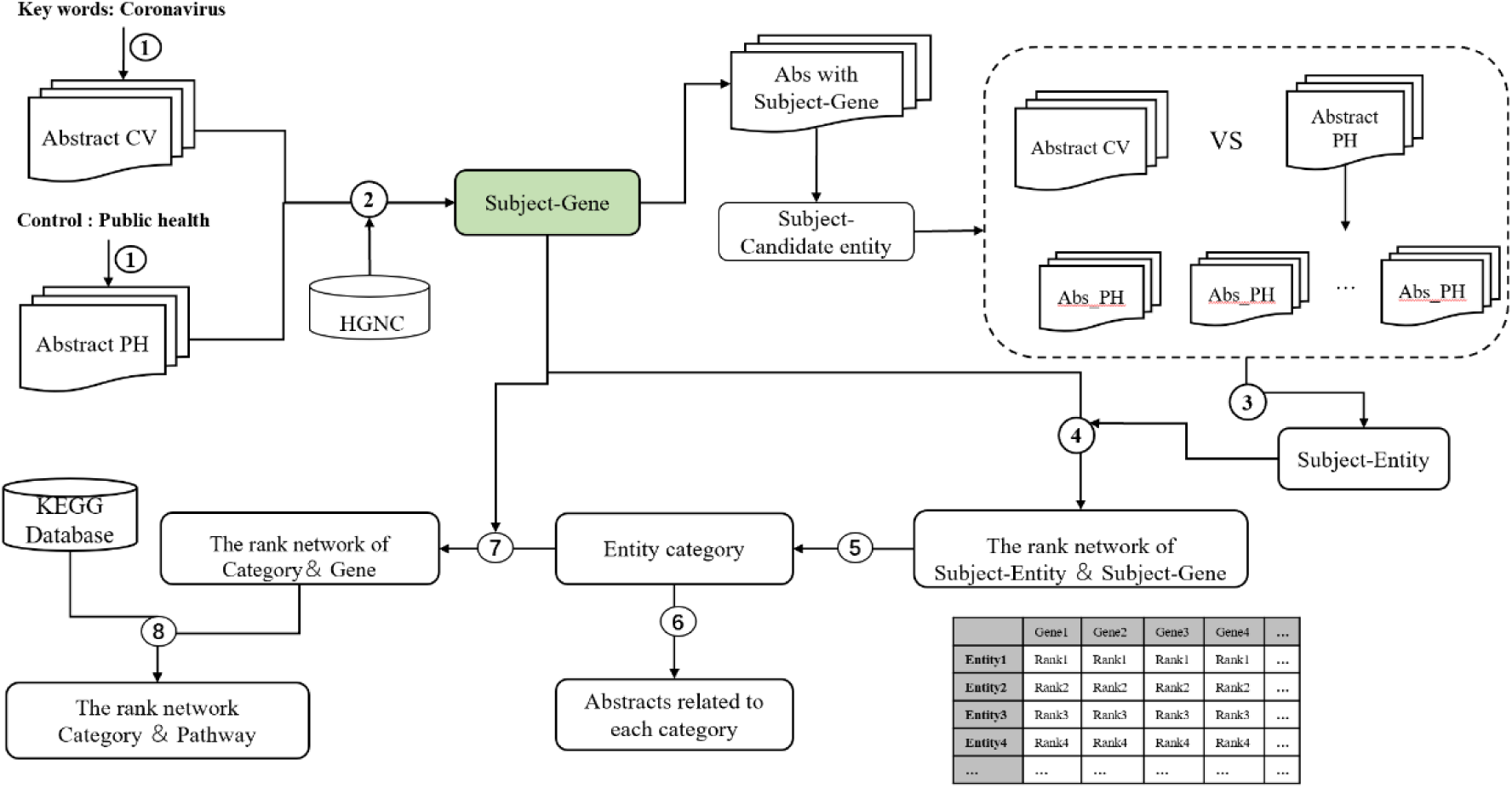
Flow chart of the knowledge-driven literature mining method.

We expect the mechanism summarized and reasoned by TWIRLS can be further supported by pathological evidence. To date, only one report of a post-mortem biopsy has been published with pathological data. Although histological examination showed bilateral diffuse alveolar damage with cellular fibromyxoid exudates, the right lung showed evidence of desquamation of pneumocytes and hyaline membrane formation, indicating acute respiratory distress syndrome (ARDS), whereas the left lung showed pulmonary edema with hyaline membrane formation, suggestive of early-phase ARDS. The pathological evidence suggests that ARDS symptoms are closely related to cytokine storm^[38]^. However, there is still a lack of histopathology-related data to support our preliminary findings generated by our machine approach.

## Methods

### Construction of the data interface

We used PubMed, the most widely used database of biological literature, as the resource for the text mining. The schematic representation of the overall study design is shown in Figure 1 and can be summarized in the following steps.

### Corpus and dictionary organization

The dataset used in this pipeline were from PubMed articles. First, PubMed was searched for articles including titles, abstracts, author and affiliation information containing the subject keyword “coronavirus”. The search results were downloaded in txt format for compiling into structured information. The text in the subject abstract set was organized and cleaned, and then assigned to specific corpuses related to coronavirus (specific corpus) and compiled into the subject dictionary. To enhance the accuracy of the effective entities associated with the keyword, we used a random corpus for comparisons. We searched for article abstracts containing the keyword “public health” and compiled the abstract set into a random corpus, and then compiled them into a randomized control dictionary, which contains a wide range of proteins, genes, and related biological entities. We also considered a balanced amount of information by setting relevant parameters to adjust the amount of text before carrying out the statistical analyses.

### Identification of genes precisely related to the subject “coronavirus”

Biological entity identification is a key step in the literature mining process. To ensure functionality of the extracted entity, we first compared the entity from the subject dictionary with the human official gene symbols in the Hugo Gene Nomenclature Commission (HGNC) database to generate subject candidate genes using standard nomenclature. In addition, the entities in the abstract were capitalized to avoid errors in the identification process. To obtain widely used gene entities that are precisely related to the subject and to determine the significance of the gene distribution in the specific texts, we calculated the difference in the distribution proportions. We searched for the subject candidate genes in the subject dictionary and the randomized control dictionary, respectively. We also counted the number of abstracts containing each subject candidate gene in each abstract set, respectively. Finally, we calculated the odds ratio of each subject candidate gene and sorted them into a list of precisely related genes (CSHG).

### Identification of all entities (CSSE) correctly related to the subject “coronavirus”

Similar to the process of identifying CSHG, we calculated whether entities were significantly distributed in a specific corpus. We counted the number of texts containing each CSHG in a specific corpus, and then counted the number of each candidate entity in the corpus subset. Next, we randomly selected the same amount of text from the random control corpus and then counted the number of each candidate entity in this subset of the random corpus. This was repeated 100-10000 times in the random corpus to generate candidate entities in the specified amount of text of the random distribution model. According to the central limit theorem (CLT), the distribution of random sampling averages of randomly distributed data always conforms to a normal distribution. Therefore, we can use the Z score to evaluate whether an entity is significant in a specific text. Here, we used a Z score cutoff value > 6.

In addition, some entities have singular and plural noun forms, and synonyms with multiple forms in the abstracts. Therefore, we numbered the subject-related entity and automatically combined nouns with plural forms and homologous words with adjectives and adverb roots into the same entity, and then assigned them the same number.

### Programming language and efficiency

Part of the algorithm was developed using the MatLab programming environment and Python language. Algorithm efficiency improvements and the targeted parallel acceleration module were developed in C/C++ language. The automated text analysis took about 4 hours to complete on a workstation with an Intel Xeon CPU E5-2690 v4 X2 (28 cores) and 128 GB of memory.

## Declarations

### Consent for publication

Not applicable.

### Ethics approval and consent to participate

Not applicable.

### Availability of data and material

All data generated or analyzed in this study are included in the published article (Table S1 and Table S2).

